# Age- and Light-Dependent Changes in the Zebrafish Olfactory Epithelium

**DOI:** 10.64898/2026.02.20.707010

**Authors:** George B. Chapman, Rania Abutarboush, Victoria Connaughton

**Affiliations:** Department of Biology, Georgetown University, Washington DC, USA; Department of Biology, Washington, DC, USA; Center for Behavioral Neuroscience, American University, Washington, DC, USA

**Keywords:** ultrastructure, light rearing level, *Danio rerio*, constant light, larvae

## Abstract

Light and transmission electron microscopy were used to identify changes in ultrastructure of the olfactory pit of larval zebrafish (*Danio rerio*) that occur as a result of age and altered environmental light levels. Larvae were reared under control/cyclic light or constant light condition until 4, 8, and 15 days postfertilization (dpf). The larval olfactory pit consisted of an epithelium that varies from simple to pseudostratified to stratified and contained three types of receptor cells: ciliated, microvillar and ciliated crypt. A variety of non-receptor cells were also identified: kinociliate non-sensory supporting cells, vesicular supporting cells, basal cells and an occasional intruder, such as a neutrophil or a lymphocyte. Microvilli projecting from microvillus receptor, kinociliate, and vesicular supporting cells were single, forked, or doubly forked. Junctional complexes were evident between a variety of cells including adjacent epidermal cells, an epidermal cell and a kinociliate cell, a kinociliate and a vesicular supporting cell, and two vesicular supporting cells. Desmosomes were also observed between adjacent cell types. With age, the olfactory epithelium thinned and vesicle number varied. In larvae reared in constant light, mitotic figures were evident, microvillar receptor cells were absent, and, at 4 dpf, some ultrastructural components were similar to those observed in 8 dpf control animals, suggesting precocious development. These findings suggest that constant light rearing alters the timing of receptor replacement, supporting previous work showing that rearing light levels impact sensory system growth and development.

## Introduction

The structure of the olfactory organ of fish has been extensively studied. Indeed, Bannister (1965) cited reports dating from 1887 in his literature review and detailed comparative study of the olfactory surface of the minnow *Phoxinus phoxinus* and the three-spined stickleback *Gasterosteus aculeatus*. As might be anticipated, the gross olfactory structures were quite different in these two species and neither one is similar to the structure of the olfactory pit of the larval zebrafish, *Danio rerio*.

However, the cells comprising the olfactory epithelium of fish (and of vertebrates in general) are remarkably similar in both the cell types present and the ultrastructural features of each type. The cells include receptor cells, supporting cells and ciliated non-sensory cells. Receptor cells can be further subdivided into three categories determined by the form of their dendritic tips: ciliated olfactory receptors, microvilli-possessing olfactory receptor cells and a previously undescribed receptor with a naked rod extending from its distal surface.^1^ The first two categories of receptor cells have been identified in almost every report on the olfactory epithelium of fish. The status of the naked rod cells remains somewhat indefinite. Later studies would add crypt receptor cells that bear both cilia and microvilli, discovered in adult zebrafish by Hansen and Zeiske (1998), and are thought to be unique to fish. Basal cells and goblet cells should probably be included, although the latter may not occur within the sensory olfactory epithelium of some fish, but are always present in surrounding non-sensory tissue. Additionally, rodlet cells, an occasional olfactory cell type^2^ also occur in the heart, gills, intestine and other organs of fish,^3^ although their origin and function are still unknown.^2^

The olfactory organ of adult zebrafish is a rosette consisting of multiple lamellae lined by the olfactory epithelium.^4,5^ Ciliated sensory neurons are found deeper in the epithelium than microvillus and crypt neurons, which occupy the surface.^2,5,6^ The zebrafish olfactory placode is a defined structure at ∼17-18 hours postfertilization (hpf)^7,8^ and axons leave the placode and enter the brain at ∼24 hpf ^5,7^. At ∼34-36 hpf, olfactory pits form by invagination of the placodes and receptor cells are evident at hatching (∼48-50 hpf).^5^ At 2.5-3 days (d) pf odor responses can be detected in the olfactory bulb^9^ though olfactory-stimulated behaviors are not reliably measured until ∼4 dpf ^9,10^ indicating functional olfactory receptor cells (and circuitry) are present at this time.^5^ As the larva continues to grow, so does the olfactory organ, with lamellae continually added throughout the life of the fish.^5^

Olfaction in fish is responsible for triggering behaviors that include finding food, reproduction, and fleeing predators. Odorant stimuli include amino acids, nucleotides, bile acids, and prostaglandins,^11^ each stimulating specific regions of the olfactory bulb.^12^ Anatomical, molecular, and physiological techniques have identified the expression of different receptor genes^7,13-15^ and/or receptor cell responses^16,17^ to these different odorant molecules in zebrafish. Specifically, both ciliated receptor cells and microvillar receptor neurons respond to amino acid stimuli. Bile acids specifically stimulate ciliated receptor cells, while nucleotides stimulate microvillar receptor neurons.^17^ The ligand for ciliated crypt receptor cells has proven to be more elusive. Both larval and adult zebrafish, like other fish, respond to odorant molecules dissolved in water^18,19^ in a pattern that suggests that different receptor types are either found on different cells or are mediated by different transduction mechanisms.^9,12,16-18,20-22^

Previous studies by ourselves and others have shown that light rearing history affects overall development of zebrafish larvae^23-25^ as well as development of the brain^26^ and retina.^23,27,28^ Interestingly, although light is not the natural stimulus for the olfactory system, circadian changes in olfactory responses have been reported in a variety of species, including *C. elegans,*^29^ *Drosophila,*^30^ and mammals.^31,32^ In particular, circadian rhythms identified in mice suggest the presence of a clock within the olfactory bulb.^31^ Though the zebrafish olfactory system is less complex than mammals^33^ and differences between species have been noted, there are also similarities.^4,12,17,34,35^ As a result, it is reasonable to hypothesize that rearing zebrafish in abnormal light conditions (i.e., constant light) may have a deleterious impact on olfactory system development.

Thus, the purpose of this study was twofold. First, we examined the developmental ultrastructure of the olfactory epithelium in larval zebrafish at 4, 8, and 15 dpf, building on previous work. Second, we determined how rearing light conditions would affect this development by rearing the larvae in constant light conditions. Given the overall deleterious effects of constant light rearing, we hypothesized that larvae reared in constant light conditions would have altered ultrastructure of their olfactory epithelium, suggesting possible functional consequences.

## Material and Methods

Adult wildtype zebrafish (*Danio rerio*) were allowed to spawn in-house and embryos were collected. At ∼24 hpf, embryos were placed into either constant light or control/cyclic light, i.e., 14 hours light/10 hours dark conditions, conditions until aged 4, 8 or 15 dpf. All embryos were maintained in fresh water at 28.5°C and fed daily from a culture of mixed *Paramecia* beginning at 4 dpf. Food density was maintained at a minimum of 10,000 *Paramecium* per liter, which is considered *ad libitum.*^36^ Standard aquarium lights (spectral range: 400-600 nm, light intensity: ∼6000-7000 cd/m^2^) were used as the light source and larvae were checked at the same time each day. All procedures were approved by American University’s Institutional Animal Care and Use Committee.

For electron microscopy, one zebrafish from each growth regimen was euthanized with 0.02% tricaine methanosulfate (MS-222) and then placed into a primary fixative consisting of 5% glutaraldehyde^37^ in double strength phosphate buffer with CaCl_2_^38^ at a pH of 7.2-7.3 for 2 hours at 4°C. All specimens were then washed four times, 15 minutes each, in buffer. They were then post-fixed in 1% osmium tetroxide in buffer at a pH of 7.2 for 2.25 hours at 4°C. Following the same washing procedure, each specimen was dehydrated through a graded (50%, 70%, 85%, 95%, 100%, 100%) ethanol-water series, followed by two changes of propylene oxide, each change taking 15 minutes. All specimens were infiltrated overnight at 20°C in a 1:1 mixture of epoxy resin (LX-112) and propylene oxide.^39^ They were further infiltrated at 20°C in a 3:1 mixture of epoxy resin and propylene oxide for 9 hours and then embedded in epoxy resin and placed into a polymerizing oven at 45°C for 24 hours, followed by another 24 hours at 60^0^C.

Thin (0.25-1.0 µm) and ultrathin (50-70 nm) sections were cut with a Sorvall Porter-Blum MT2-B ultramicrotome and a Delaware diamond knife. The thin sections were mounted on glass slides and stained with an aqueous mixture of 1% methylene blue, 1% azure A and 1% sodium borate.^40^ Photomicrographs were taken with a Bausch and Lomb research microscope equipped with Nikon lenses and No. 4 Ilex Optical Company camera. The ultrathin sections were collected on uncoated 200, 300, or 400 mesh copper grids, stained with uranyl acetate in 50% ethanol for 20 minutes^41^ and 0.4% lead citrate for 10 minutes.^42^ Electron microscopy was conducted with a JEOL JEM-1010 transmission electron microscope. Either Kodak Ektapan 3162 or Kodak T-MAX 100 film was used for photomicrographs and Kodak 4489 electron microscope film was used for electron micrographs. Enlarged prints were made on either Ilford Ilfospeed RC Deluxe 2 paper or Kodak Kodabrome 2 RC paper with a Simmon Bros. Inc Omega Model D2 photographic enlarger. A minimum of 10 thin (0.25-1.0 µm) and 5-10 ultrathin (50-70 nm) sections, each including several hundred cells, were examined for each age and light regimen.

## Results

### Development of the olfactory epithelium at 4, 8, and 15 dpf

#### 4 dpf

At this age, the diameter of the olfactory pit opening measures ∼ 70 µm (**Fig. 1a**). In a higher magnification image closer to the anterior rim of the olfactory pit (**Fig. 1b**), the diameter of the olfactory pit opening is smaller, ∼ 25 µm. Further examination of the luminal surface of the olfactory pit (**Fig. 1b**, moving from left to right) revealed additional structural components. For example, at the left edge and bottom center, the previously undisclosed openings of vesicles (from vesicular supporting or VS cells) into the lumen of the olfactory pit appear. Many knobs of ciliated receptor (CR) cells are prominent as are the irregularly shaped microvilli of microvillar receptor (MR) cells. A very rarely seen ciliated crypt receptor (CCR) cell occurs at the bottom center of the figure. At the top right edge, two non-sensory kinociliate supporting (KS) cells with their broad surface and many cilia appear at the junction with the epidermal cell; one to five of these KS cells have been seen at such sites. KS cells also possess microvilli. VS cells surround CR cells. At higher magnification (**Fig. 2a**), a KS cell with cilia, striated ciliary rootlets, and microvilli appears at the left side of a VS cell.

**Fig. 1.**
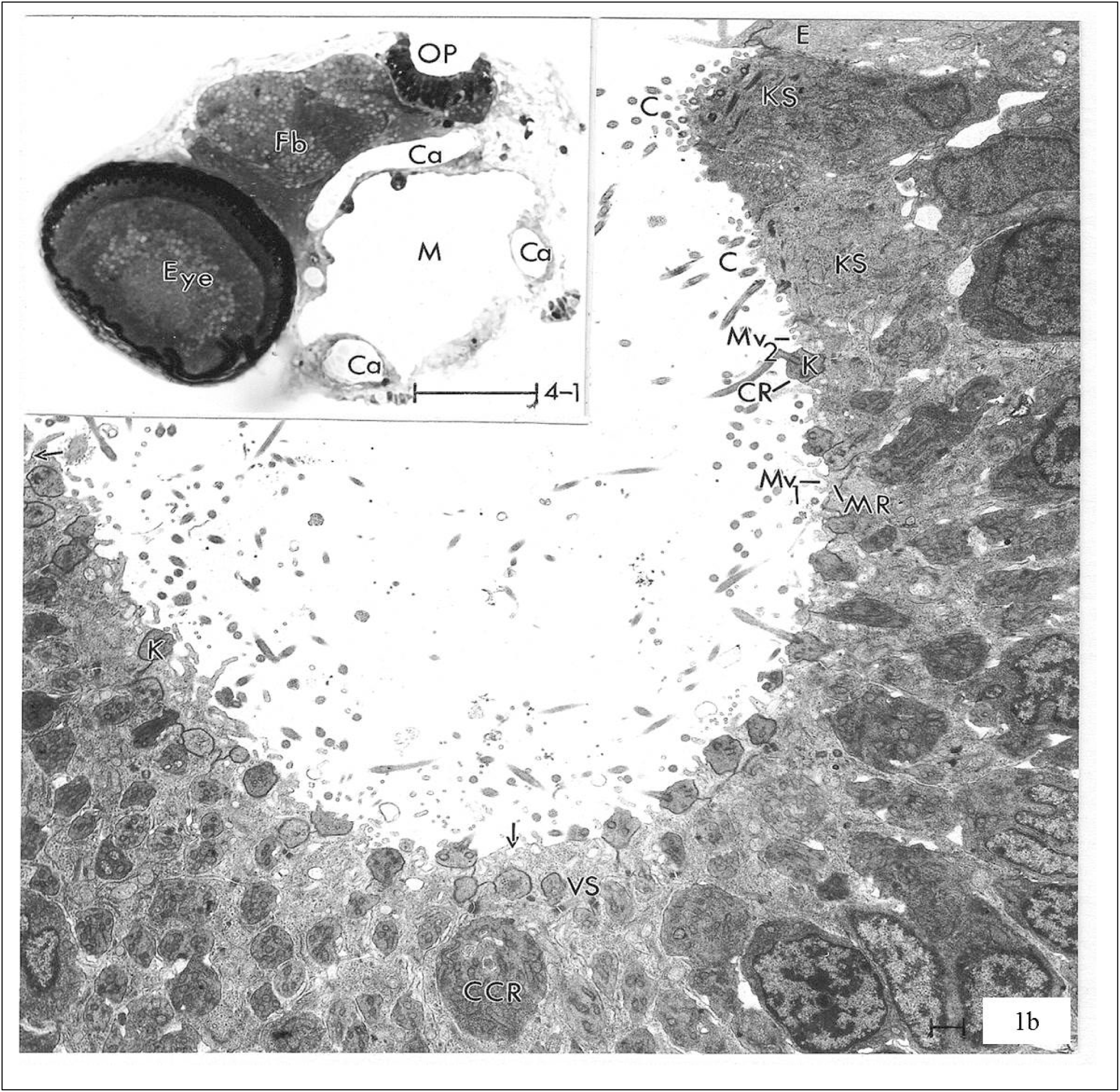
4 dpf control larvae. **(a)** Head of a 4 dpf larval zebrafish reared in control conditions. The plane of section permits showing only one eye (Eye) and one olfactory pit (OP). Fb, forebrain; Ca, cartilage plate; M, mouth. **(b)** Higher magnification, vertical section of the olfactory pit shows major features of interest, counterclockwise from the left edge: the opening of a vesicle (arrow) of a VS cell, a knob (K) of a CR cell, another opening of a vesicle (arrow) of a VS, a CCR cell, a MR cell and its projecting microvilli (Mv_1_), a CR with a knob (K) that has a projecting cilium, microvilli (Mv_2_) projecting from a KS cell, cilia (C) projecting from the KS cell, another KS cell with cilia (C) and an epithelial cell (E) in the skin. Scale bar in **(a)** = 100 µm. Scale bar in **(b)** = 1µm. VS = vesicular support; KS = kinociliate support; CCR = ciliated crypt; CR = ciliated receptor.

**Fig. 2.**
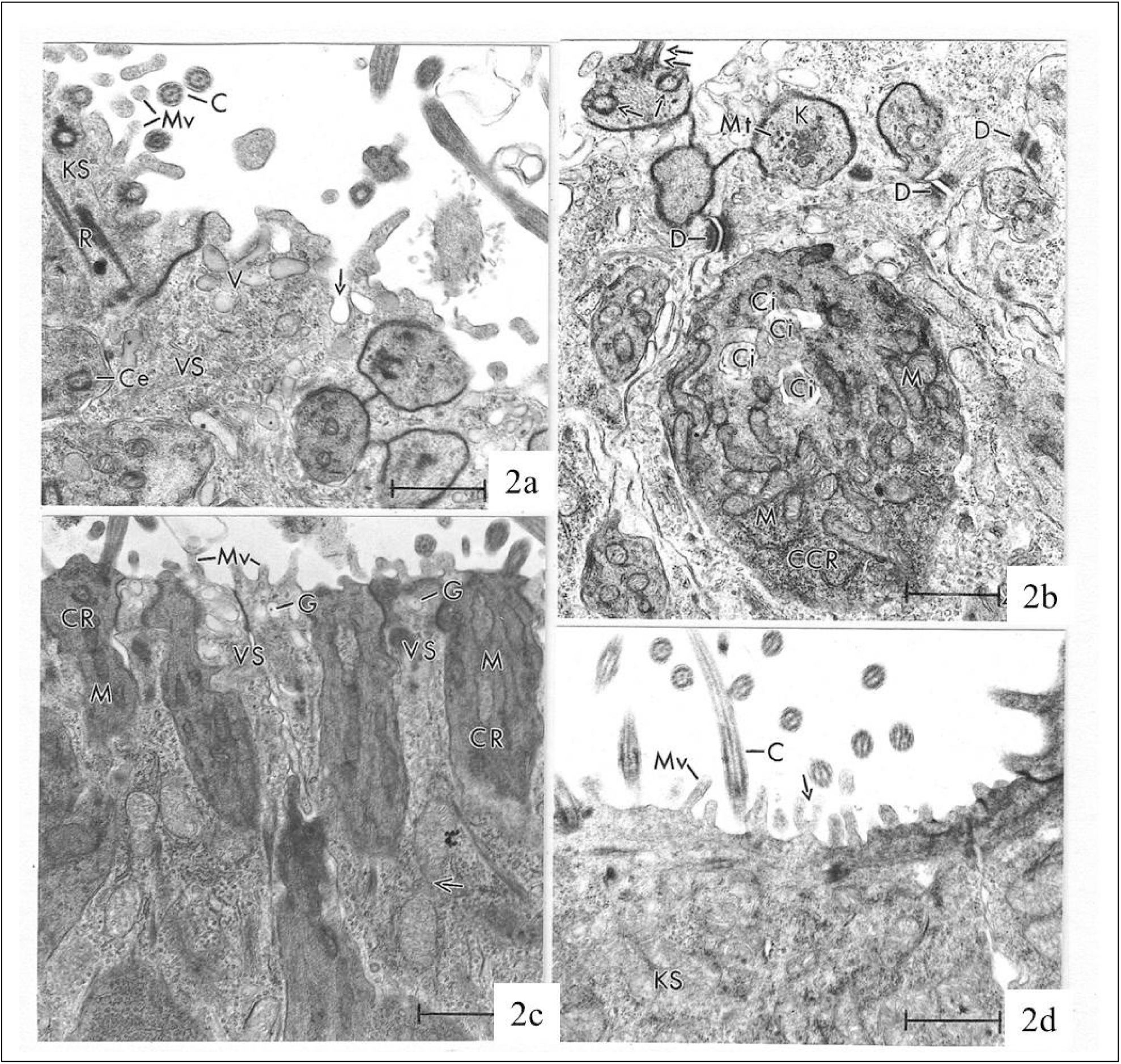
4 dpf control larvae. **(a)** High magnification micrograph of the upper left edge of Fig. 1b shows details of an opening vesicle (arrow) and vesicles (V) of a vesicular supporting cell. A centriole (Ce) of an adjacent ciliated receptor cell knob, and striated ciliary rootlet (R), microvilli (Mv) and cilia (C) of a KS cell are also present. **(b)** Higher magnification of the CCR cell of Fig. 1b shows its four in-sunk cilia (Ci) and many large mitochondria (M). Three desmosomes (D) between surrounding VS, several microtubules (Mt) in the knob (K) of ciliated receptor cell and another such knob with one longitudinally (double arrow) sectioned base of a cilium and two transversely (arrow) sectioned basal bodies are present. **(c)** A region of the olfactory surface with a high concentration of ciliated receptor cells (CR), each containing several mitochondria (M). Several VS cells surround the receptor cells. Each labeled VS cells contains a vesicle with a granule (G). All VS cells have microvilli (Mv) and the VS cell on the right contains a dividing mitochondrion (arrow). **(d)** A portion of the broad surface of a KS cell showing simple microvilli (Mv), a forked microvillus (arrow) and a cilium (C). All scale bars = 1 µm. VS = vesicular support; KS = kinociliate support; CCR = ciliated crypt.

The latter contains several vesicles with finely granular, low electron density content and a distinct limiting membrane that appears identical to the membrane that is confluent with the plasma membrane of the site of a vesicle’s opening to the pit lumen. The omega-like appearance of these vesicles suggests exocytotic release of vesicular content onto the luminal surface of the pit.

Ciliary rootlets are separated by ∼ 70 nm and their diameter is ∼ 140 nm at 4 dpf. At higher magnification, the CCR cell (**Fig. 2b**) reveals four “in-sunk” cilia^5^ and numerous mitochondria. The figure also includes the knobs of seven CR cells and three desmosomal junctions between VS cells. The knob at the top center contains a group of microtubules, an observation unique to this treatment group. The preferred disposition of VS cells is around CR cells (**Fig. 2c**). Also evident were short microvilli projecting from the apical surface of VS cells and electron-lucent vesicles containing dense granules. The largest vesicle presenting a circular profile was ∼ 200 nm in diameter and the largest granule was ∼ 40 nm in diameter.

The broad surface of KS cells with both simple and forked microvilli and cilia (**Fig. 2d**) is rarely encountered. The rarity of that cell type in the 4 dpf group may be age-related. At the junction between the non-olfactory (skin) epithelium, with its numerous projections, and the KS cells, several ultrastructural details become obvious (**Fig. 3a, b**), including (1) projections from the surface of the non-olfactory epithelial cells; (2) desmosomal junctions between the non-olfactory and KS cells and between KS cells and VS cells; (3) the preferential orientation of the cilia of KS cells, with a plane through each central pair of microtubules in the cilia in a vast field of cilia having the same direction;^6^ (4) short microvilli and long cilia projecting from the same KS cell, and (5) ciliary rootlets with nearly identical angulation. Some of the same features, e.g., preferential ciliary orientation, are also present on the opposite side of the rim of the olfactory pit (**Fig. 3c**) indicating the expanse over which they may extend and suggesting that KS cells tend to be arranged as a ring around the periphery/perimeter of the olfactory pit (though they can also occur in other portions of the olfactory pits).

**Fig. 3.**
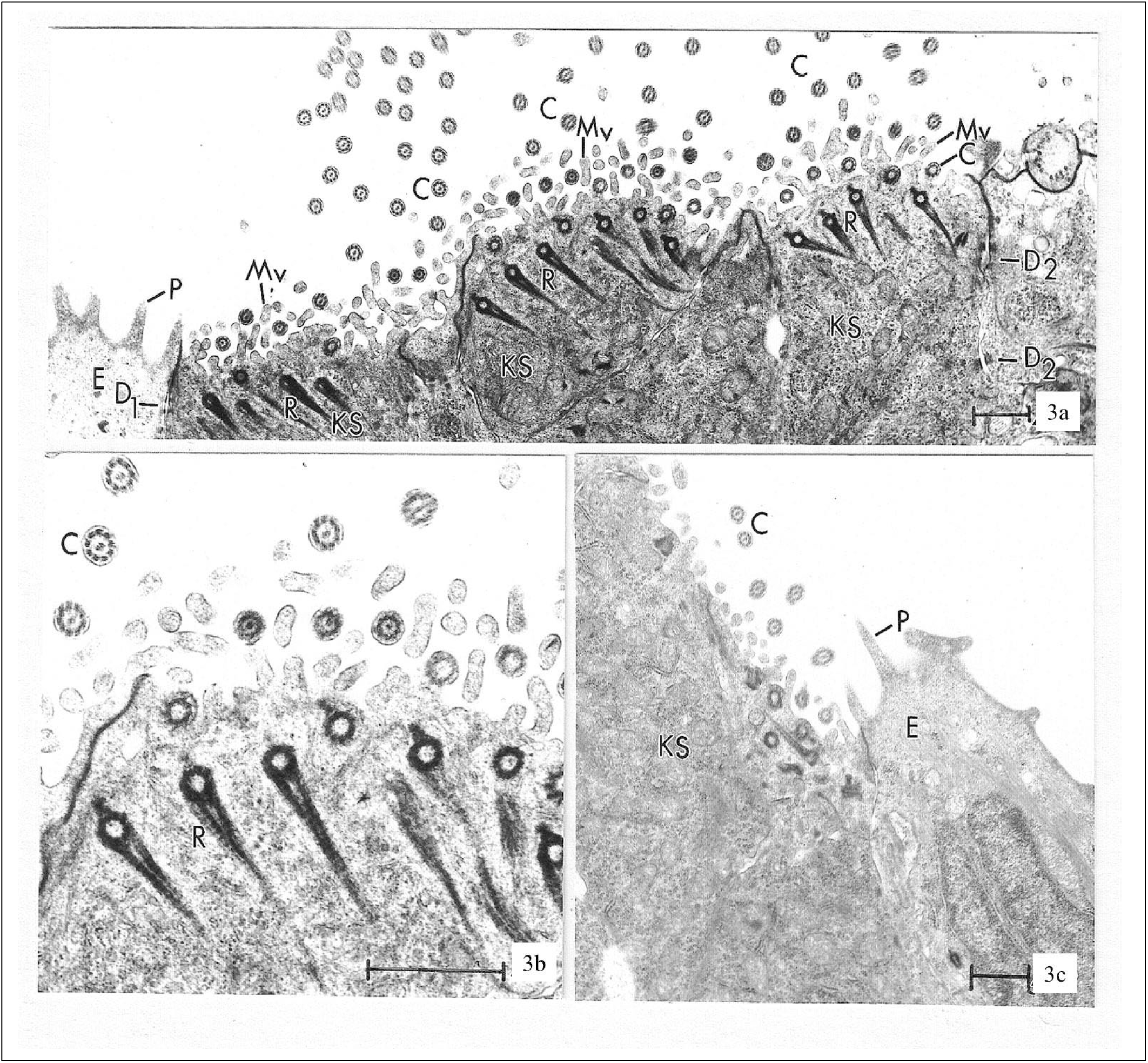
4 dpf control larvae. **(a)** At the junction between the skin and the olfactory epithelium, there are three KS cells, each containing several striated ciliary rootlets (R), microvilli (Mv) and cilia (C). E, epidermal cell in skin; P, projection from skin; D_1_, desmosome between epidermal cell and kinociliate supporting cell; D_2_, desmosome between VS and KS cells. All cilia reveal the same preferential orientation of the central doublet microtubules of the axoneme. All ciliary rootlets display the same directionality. **(b)** Higher magnification to render clearer the features in panel *A*. **(c)** Junction between the epidermis of the skin (E), with projections (P), and a KS cell. This junction is on the opposite side of the olfactory pit from the junction in panel *A*. The preferential orientation of the central doublet microtubules of the axoneme of the cilia (C) appears again. All scale bars = 1 µm. VS = vesicular support; KS = kinociliate support.

In sum, the olfactory pit ultrastructure at 4 dpf includes (1) a single CCR cell, (2) vesicle membrane confluence with the plasma membrane at a site of exocytosis into the olfactory pit lumen, (3) ciliary rootlet periodicity/spacing, (4) vesicle and vesicular granule sizes, (5) forked microvilli in KS cells, and (6) microtubules in the knob of a CR cell (**Table 1**).

**Table 1.** Summary of notable observations. List of structures and their identification in the olfactory pit/epithelium of zebrafish larvae aged 4, 8, and 15 days postfertilization (dpf) reared under control (cyclic) light or constant light (italics) conditions. Specific figures where the different structures can be observed are given at the right.

| structure | 4 dpf |  | 8 dpf |  | 15 dpf | Figure(s) |
| --- | --- | --- | --- | --- | --- | --- |
|  | control | *Constant light* | control | *Constant light* | control |  |
| Ciliated Receptor Cell (CR) | X | *X* | X | *X* | X | 1b; 2c; 4a, c; 7c; *11a, b; 12a; 13* |
| Microvillar Receptor Cell (MR) | X |  | X |  | X | 1b; 4a-c; 5a, b; 6; 8b |
| Ciliated Crypt Receptor Cell (CCR) | X | *X* | X | *X* |  | 1b; 4a, b; *12a, b; 14* |
| Kinociliate Supporting Cell (KS), rim | X | *X* | X |  | X | 3a,c; 4a; 5a,c,d; 6; *10b* |
| KS cell, central |  | *X* | X | *X* | X | 5a,b; 6; *11b; 12a; 13* |
| Vesicular Supporting Cell (VS) | X | *X* | X | *X* | X | 1b; 4c; 6; 7b,c; *12a; 13* |
| Dark Basal Cell |  |  |  | *X* |  | *13* |
| Light Basal Cell |  |  |  | *X* |  | *13* |
| Mitotic Figures |  | *X* |  |  |  | *10c,d; 11a* |
| Neutrophils |  |  |  | *X* |  | *13* |
| Lymphocytes |  |  | X |  |  | 4b |
| Monocilium |  |  | X |  |  | 5c |
| Axon Bundle |  |  |  |  | X | 6 |
| Axon Singles from KS |  |  |  |  | X | 8a; 9a |
| Axon Singles from non-KS |  |  |  |  | X | 9a |
| Axon singles from probably MR and CR cells |  |  |  |  | X | 6 |
| Microvilli of VS | X |  |  |  | X | 2c; 6; 7b,c |
| Microvilli of MR | X |  | X |  |  | 1b; 4c; 5a |
| Microvilli of KS | X | *X* | X | *X* | X | 1b; 4c; 8a; *12a; 13* |
| Forked Microvilli of MR |  |  | X |  | X | 4a; 6; 8b |
| Forked Microvilli of KS | X |  | X |  | X | 2d; 4a,c; 5c; 9a |
| Doubly-forked microvillus |  |  |  |  | X | 8a |
| Vesicles of VS | X | *X* |  |  | X | 2a; 6; 7b,c; 8c,d; *12a,b* |
| Vesicles open to olfactory pit lumen | X | *X* | X |  |  | 1b; 2a; 4b; *12a,b* |
| Vesicle granule | X | *X* |  |  |  | 2c; *12a,b* |
| Knob of CR | X | *X* | X | *X* | X | 1b; 4a; 5a; 6; 7c; *11b; 12a; 13; 14* |
| Microtubules in knob of CR | X |  |  |  |  | 2b |
| Preferential orientation of axonemal central doublet | X | *X* |  |  | X | 3a-c; 9b |
| Door knob-like projections |  |  |  | *X* | X | 6; 7b; *13* |
| Structure-less projection |  |  | X |  | X | 5a; 7c |
| Electron-dense mitochondria in KS |  |  |  |  | X | 7a |
| Electron-lucent mitochondria in VS | X |  | X |  | X | 5b; 7a |
| Electron-dense mitochondria in MR |  |  | X |  |  | 65 |
| Rootlets of Cilia | X |  | X |  |  | 2a; 3a,b; 5c |
| Epidermal Cell and Projection | X |  | X |  |  | 3a,c; 4a; 5a,c,d; *10b* |
| Circular Profile Groups |  | *X* |  |  |  | *11c* |

#### 8 dpf

The olfactory epithelium at 8 dpf includes all structures shown at 4 dpf, except that at 8 dpf forked microvilli now project from both MR and KS cells (**Fig. 4a**). Several CR cells appear, the left one displaying its nucleus, and the right one displaying its knob, with a projecting cilium (**Fig. 4a**). Cilia from other CR cell knobs are also shown. The presence of a portion of an epidermal cell, with projections (top left and top right edges in **Fig.4a**), at the junction of the epidermis with the KS cells, indicates that the KS cells are dispersed in a circle around the perimeter of the rim of the olfactory pit. Figure 4a clearly illustrates the difference between the broad surface of the KS cells and the knob-like apical terminations of the ciliated dendritic processes of the CR cells.

**Fig. 4.**
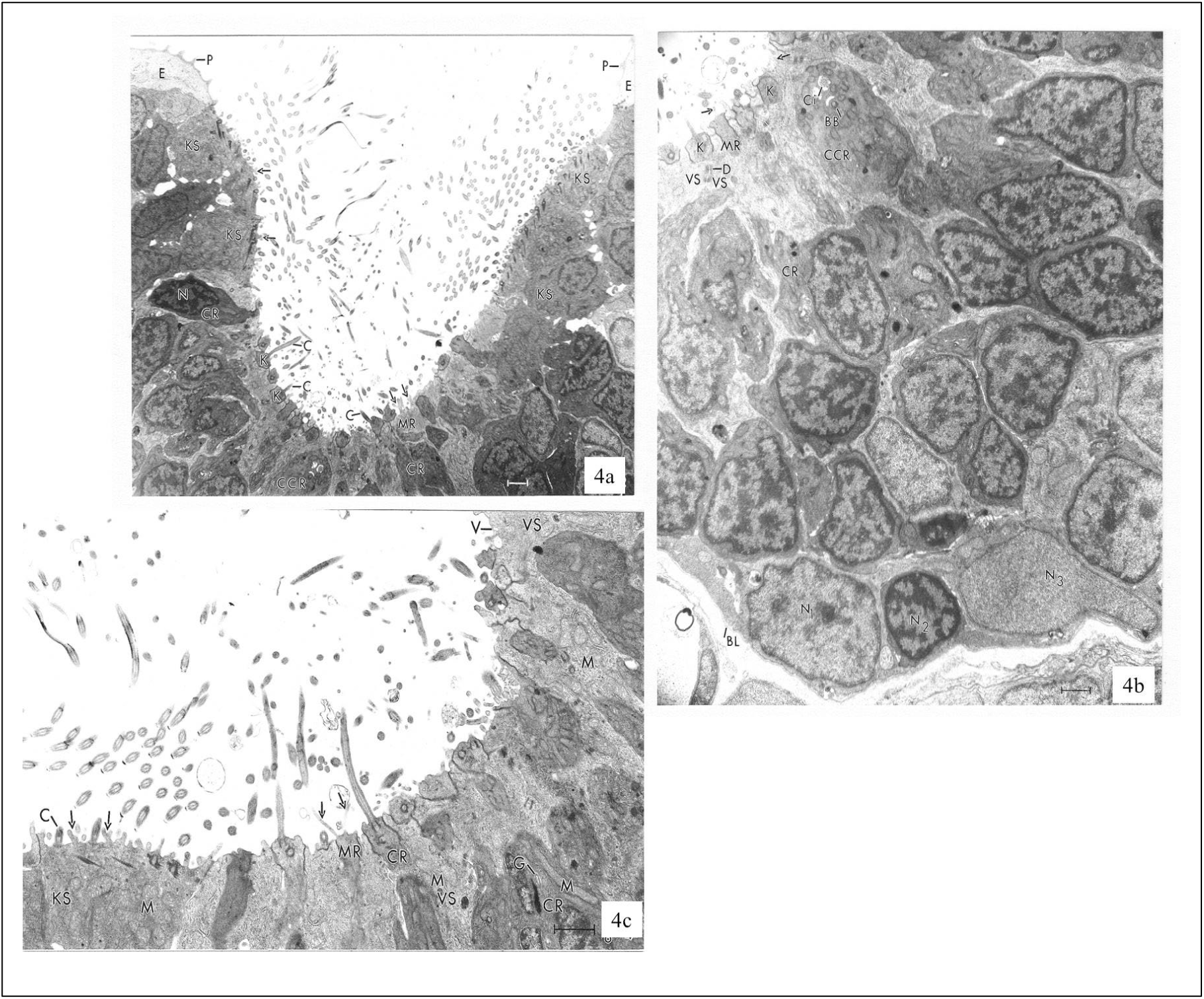
8 dpf control larvae. **(a)** Vertical micrograph of a section of olfactory pit at 8 dpf. Skin epidermal cells (E) with their projections (P) are at the top left and right edges. The position of the several KS cells relative to the skin cells indicates that these cells form a ring around the perimeter of the pit opening. Several forked microvilli (arrows) extend from the KS cells on the left side and two forked microvilli (arrows) extend from a MR cell near the bottom of the olfactory pit. A CR cell with a nucleus (N) is at the left and another CR without a nucleus showing, but with a cilium (C) projecting from its knob is at the bottom. Two ciliated knobs (K) are just below the CR with the nucleus. A CCR cell is at the bottom center of the figure. **(b)** Higher magnification micrograph shows a MR cell with a microvillus (arrow), knobs (K) of two CR cells, a desmosome (D) between two VS cells, a vesicle (arrowhead) opening to the lumen of the olfactory pit, a CCR cell with an in-sunk cilium (Ci) and a basal body (BB). Three different nuclear appearances in basal cells are obvious. One nucleus (N_1_) contains a small amount of heterochromatin within its slightly irregular envelope. Another nucleus (N_2_) occupies most of the cell volume and contains abundant heterochromatin (the cell is believed to be a lymphocyte). The third nucleus (N_3_) reveals no heterochromatin. BL, basal lamina. **(c)** Slightly higher magnification of an adjacent section. The KS cell at the left reveals cilia (C), forked microvilli (arrow), single microvilli (double-arrow), and numerous apically located mitochondria (M). The MR cell near the center has two slender, single microvilli (arrow). The CR cell to its right shows a rarely attained long portion of a cilium (C) as well as its basal body (BB). The VS cell to its right contains numerous apically located mitochondria (M). The CR cell to its right contains a Golgi body (G) and an unusually long mitochondrion (M). The VS cell at the extreme right contains numerous apically located mitochondria (M), a small vesicle (V_1_) and a larger vesicle (V_2_) apparently opening to the luminal surface. All scale bars = 1 µm. KS = kinociliate support; MR = microvillar receptor; CR = ciliated receptor; CCR = ciliated crypt receptor.

A section (**Fig. 4b**) including the entire thickness of the olfactory pit epithelium permits analysis of the nucleoplasm and its component cells. Along the basal lamina, the basal cells display three distinctly different nucleoplasmic appearances. The first is low in electron density with patches of two different density heterochromatin scattered within it and a thin layer of less dense heterochromatin disposed along the inner surface of the slightly contoured nuclear envelope. This cell is elongated horizontally and has extensions along the basal lamina. The second nucleoplasm comprises a smaller proportion of the nuclear content and has patches of one very electron dense heterochromatin in the central portion of the nucleus and disposed along the inner surface of the essentially contour-less nuclear envelope. The cell appears smaller than the other two types and round in section. Only one of these cells was seen. The third nucleoplasm is distinctly homogeneous in appearance and is essentially devoid of any heterochromatin. The cell is horizontally elongated, similar to the first type. It is reasonable to think that the first and third cell types may be precursors of most or all of the apical cells. One CCR cell is in the field and several knobs of CR cells are also present. VS cells joined by desmosomes are present, but the only vesicle noted, with one opening to the lumen of the pit, is obscure. Vesicle formation may occur irregularly in these cells.

Forked microvilli were rarely seen at 8 dpf, so a KS cell with two forked microvilli, cilia, and many large mitochondria is notable (**Fig. 4c**). The figure also includes the very rarely seen dendritic end of an MR cell that contains two microvilli, and a portion of a CR cell with a very long mitochondrion and what is probably a Golgi body (identified on the basis of its appreciable electron density). VS cells have the lowest cytoplasmic electron density and contain abundant but low electron dense mitochondria and few vesicles – a puzzling finding given that vesicles were very rarely seen within these cells at 8 dpf compared to 4 dpf.

The junction between the skin and olfactory pit, including opposite sides of the rim of the pit (**Fig. 5a**), presents several interesting structural features. At the left side of the figure, there are five KS cells at the junction with the skin. At the right side, there are four KS cells while near the bottom of the olfactory pit, there is only one. Two MR cells appear at the depth of the olfactory pit, but microvilli were only observed on the one on the right. The many knobs of the CR cells indicate that CR cells greatly outnumber the MR cells. The combination of structural features revealed in the survey figure (**Fig. 5a**) is rare and presented at higher magnification (**Fig. 5b**). The two MR cells show similarly appearing nuclei. The MR cell on the left now clearly shows its 3 µm long electron-dense mitochondrion; the one on the right more clearly shows its microvilli. The VS cell on the left is devoid of vesicles and has a prominent long mitochondrion. It is interesting that the mitochondrion in the MR cell has a much greater electron density than the mitochondrion in the VS cell, a difference shared by their cytoplasmic electron densities. One VS cell presents a projection seen only once at 4 dpf.

**Fig. 5.**
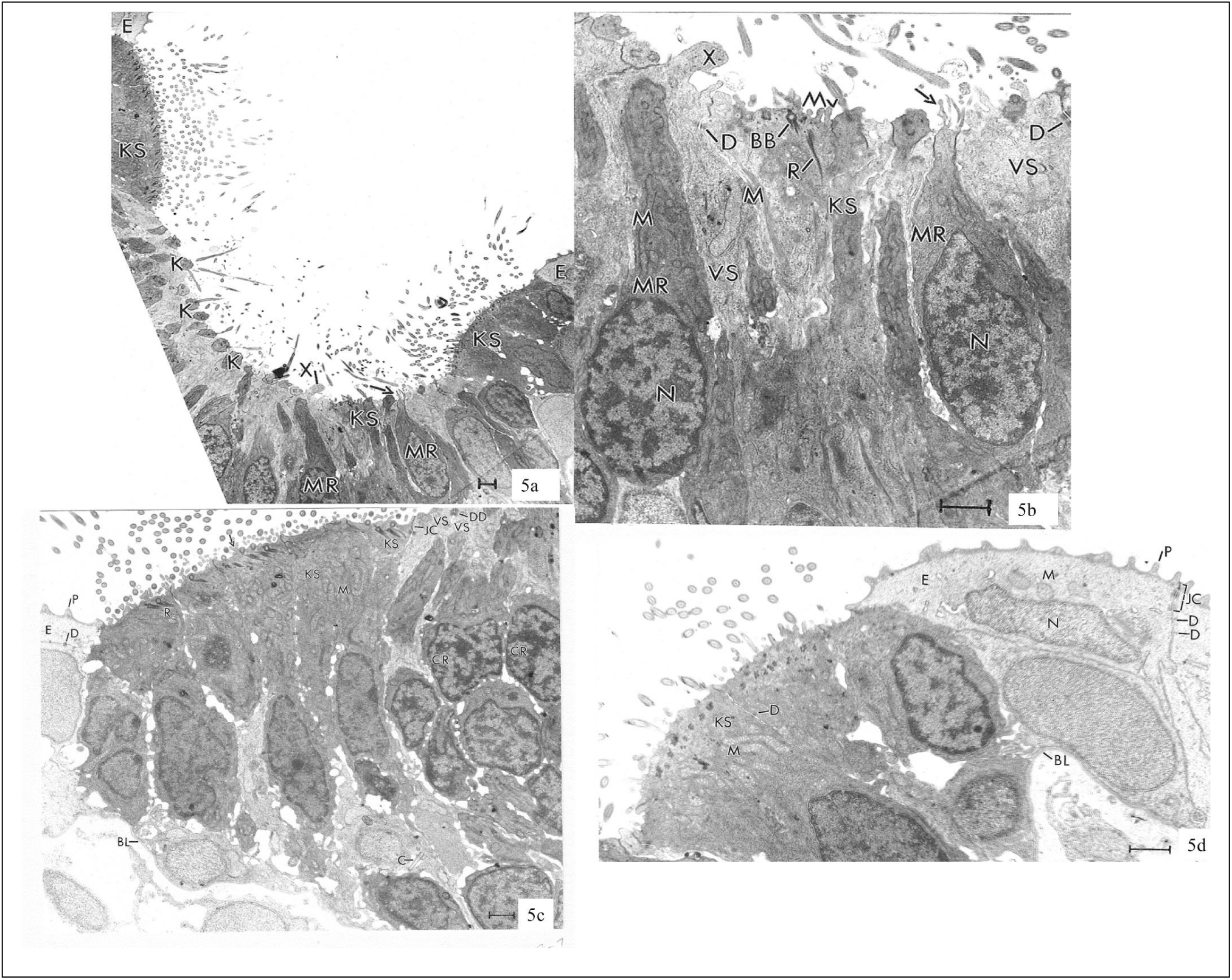
8 dpf control larvae. **(a)** This low magnification survey figure discloses KS cells occurring as a group at the junction with the skin epidermis (E) at each side, their usual location, but also shows a lone KS cell near the bottom of the olfactory pit. On the left side of the micrograph, one knob (K) of a ciliated receptor cell has three cilia, another knob has only one cilium, and a third knob has no cilium (at least in this plane of section). Several MR cells are present, but only the one to the right discloses microvilli (arrow). A unique structureless projection (X) from the luminal surface appears at the bottom of the olfactory pit. **(b)** At higher magnification, the unique structureless projection (X) reveals no organelles. The mitochondria (M) in the MR cells are more electron-dense than those in the VS cells. One MR cell has several microvilli (arrow). N, nucleus. The lone KS cell reveals its basal bodies (BB), microvilli (Mv) and striated rootlets (R). Several VS cells display desmososmes (D). N, nucleus of MR cells. **(c)** The junction between the skin epidermis (E), with its projections (P) and a diminutive desmosome (D), the usual group of several KS cells, with striated ciliary rootlets (R) a forked microvillus (arrow), electron-dense mitochondria (M) and a junctional complex (JC) with a VS cell, appear. DD, double desmosome between two VS cells; BL, basal lamina; C, cilium of monociliate cell. **(d)** An enlargement of the right side of panel *a* reveals that the skin epidermal cell (E) displays many projections (P), a junctional complex (JC), and several desmosomes (D) between like cells, several mitochondria (M) and a nucleus (N). The KS cells contain diminutive desmosomes (D) between like cells and elongate mitochondria (M). Bl, basal lamina. All scale bars = 1 µm. KS = kinociliate support; MR = microvillar receptor; CR = ciliated receptor.

The left edge of the olfactory pit (**Fig. 5c**) includes epithelial cells that border the pit and display their projections and desmosomal connections to each other. As noted previously, they contain large regions devoid of mitochondria. The KS cells have many mitochondria, ciliary rootlets, forked microvilli, and a nucleoplasm that is essentially devoid of the heterochromatin seen in the nuclei of the CR cells located deeper into the olfactory pit (to the right in the micrograph). Where the angle of section is appropriate, the basal lamina beneath the olfactory epithelium is apparent. Perhaps the most intriguing cytological feature of this field is the isolated cilium projecting toward the basal lamina. This was the only such cilium seen. Also present is a double desmosome (= two closely associated desmosomes) between two VS cells and a junctional complex between a KS cell and a VS cell.

The right edge of the olfactory pit (**Fig. 5d**) presents several of the same features shown on the left, as well as several new features. The elongate supranuclear mitochondria of the KS cells are evident as are desmosomes between two KS cells. The nucleus of an epithelial cell in the surface layer of stratified squamous epithelium bordering the olfactory pit can be identified as can three mitochondria in the cytoplasm of that cell. A junctional complex and two very small desmosomes appear between two of the surface epithelial cells. The greater extent of skin surface includes a greater number of projections from the surface epithelial cell thereby giving a more accurate indication of the number of such projections (∼ 4 per µm^2^). The basal lamina is clearly continuous beneath the skin and the olfactory epithelium.

In sum, the olfactory epithelium at 8 dpf includes (1) forked microvilli projecting from *both* MR and KS cells, (2) basal cell differences, (3), basal lamina, (4) CCR cell, (5) microvilli of MR cells, (6) mitochondrial differences in size and electron density, (7) rarity of vesicles, (8) a monocilium (unique to 8 dpf), and (9) a lone KS cell near the center of an olfactory pit (**Table 1**).

#### 15 dpf

At 15dpf, the two olfactory bulbs project rostrally from the forebrain and the olfactory epithelium is thinner than that observed at either 4 or 8 dpf. The luminal surface at this age (**Fig. 6**) presents several especially interesting structural features. An MR cell with a forked microvillus is present as is another MR cell with several microvilli positioned several cells to the right of the previous one, providing a rare opportunity to display the entire cell body except for the axon. A single KS cell is located distant from the rim of the olfactory pit and the several KS cells. A VS cell with a single microvillus and several vesicles is centrally located. Also evident is a short expanse of basal surface characterized by doorknob-like projections that extend from basal cells toward the basal lamina which covers them. These were not seen in the younger larvae. A rarely seen bundle of axons appears near the basal edge of the epithelium. Equally rarely seen are what appear to be axons projecting from several cells considered to be either KS or MR cells because of the electron density of their nuclei and cytoplasm. The connective tissue underlying the olfactory epithelium at this age contains several fibroblasts with long, slender cytoplasmic extensions. Very few knobs of CR cells appear in this area.

**Fig. 6.**
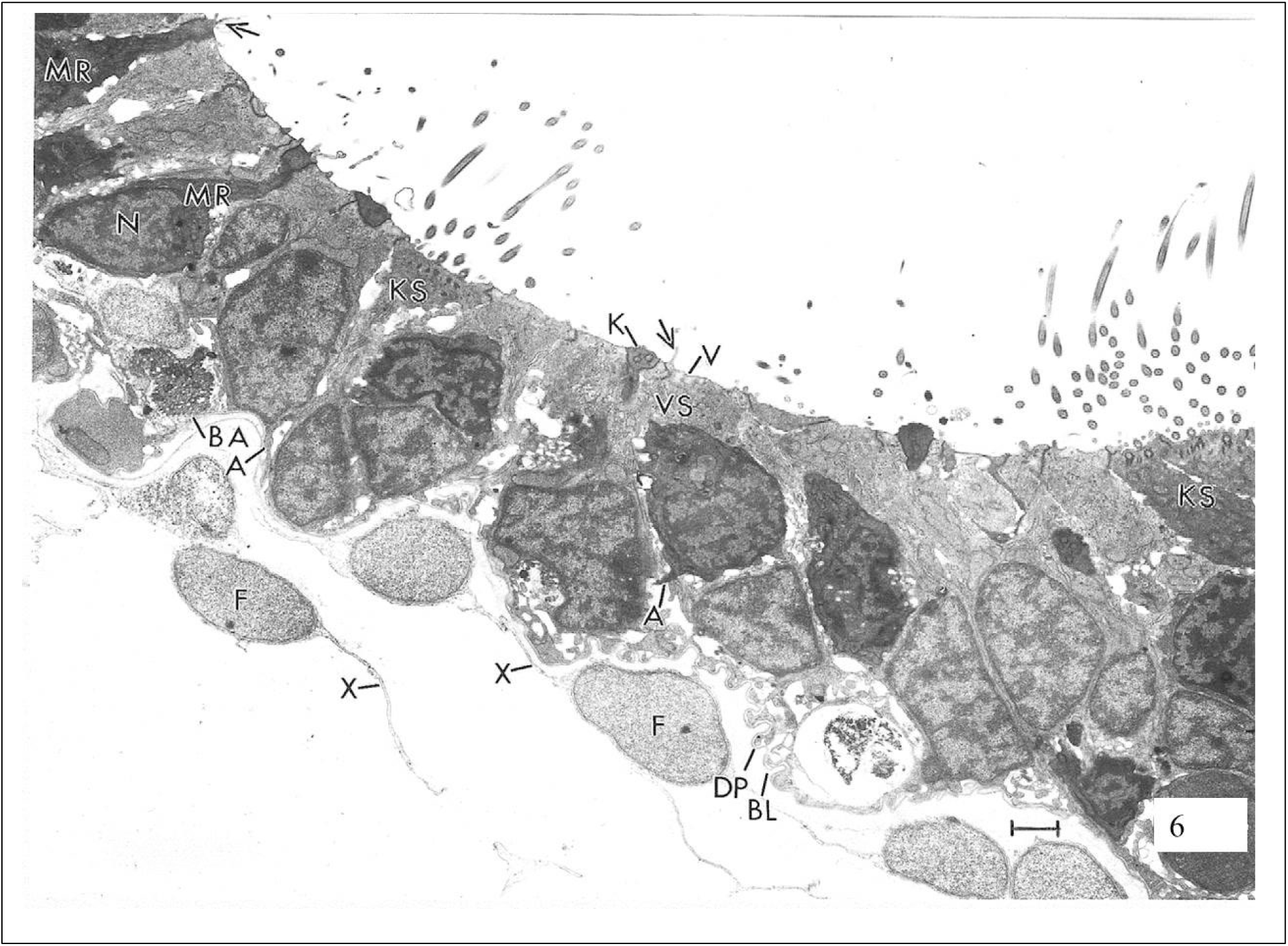
15 dpf control larvae. At 15 dpf, the olfactory pit epithelium is thinner and the most notable structures along the luminal surface include from the left, the MR cell with forked microvilli (arrow), an MR cell with its nucleus (N), a single KS cell distant from the olfactory pit rim and extending through the total thickness of the epithelium, a knob (K) of a CR cell, a microvillus (arrow) of a VS cell with vesicles (V), and a group of several KS cells. Several (probably MR) cells display axonal extensions (A). Several rarely seen door-knob like projections (DP) and a bundle of axons (BA) are positioned at the basal lamina (BL). Several fibroblasts (F) display their long, slender processes (X). Scales bar = 1 µm. KS = kinociliate support; MR = microvillar receptor; CR = ciliated receptor; VS = vesicular support.

There was large variation in cell types and in their mitochondria and nuclei within the olfactory epithelium at 15 dpf. An example of this variation is seen in the mitochondria within the different cell types (**Fig.7a**). In KS cells, the mitochondria were located supranuclearly and they were elongated and very electron dense like the cytoplasm of the cell. In VS cells, the mitochondria tend to be similar in location and shape but lower in electron density and there is often a roundish mitochondrion located either at the base of the nucleus or at the very base of the cell. At 4 and 8 dpf the VS cells also consistently presented mitochondria of lower electron density. The nuclei also present several different appearances. Some are very electron-dense and contain abundant dense heterochromatin, while others are less dense in both of those features. A third type has a homogeneous appearance, and a fourth type appears to be the smallest and is possessed of a small quantity of heterochromatin. Fibroblast nuclei, though larger, tend to resemble the fourth type.

VS cells, even when prepared identically, preserve somewhat differently. Some (**Fig. 7b**) are quite well preserved and show apically located vesicles with a limiting membrane and a very low electron density content. These VS cells also have mitochondria with low electron density (and a tendency to divide) and a few short microvilli and junctional complex contacts to others. Several doorknob-like projections extend from the KS cell at the junction with the skin. Although of slightly lower preservation quality, **Fig. 7c** is included because it provides information such as length of a microvillus and vesicles with a slightly more electron-dense content. The structureless projection (**Fig. 7c**, right edge) was seen only at 15 dpf.

**Fig. 7.**
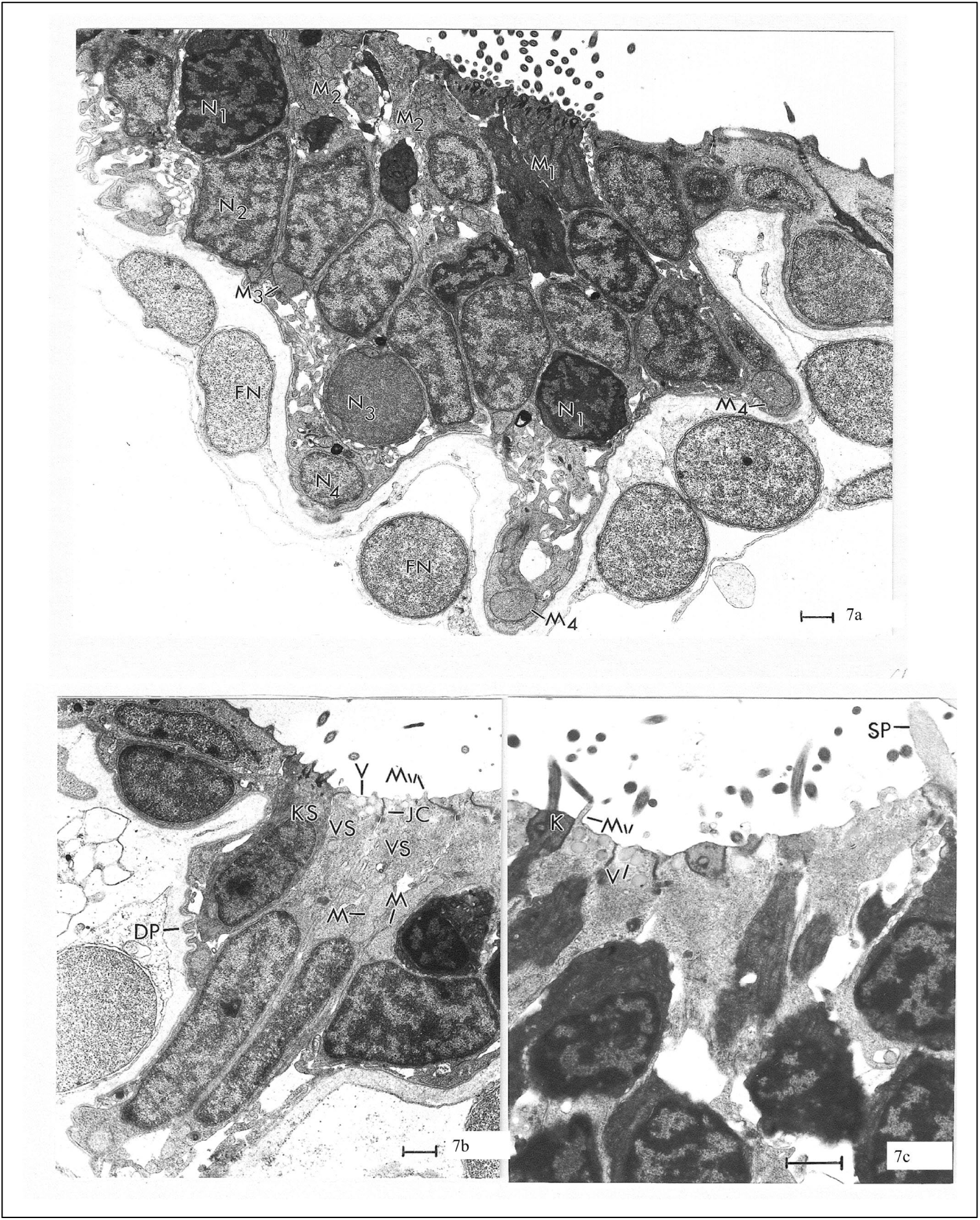
15 dpf control larvae. **(a)** There is a striking variation in mitochondria in 15 dpf control larvae. In the KS cells, the mitochondria (M_1_) tend to be concentrated supranuclearly and are elongate and electron-dense. In VS cells, the mitochondria (M_2_) tend to be concentrated supranuclearly and are elongate but are less electron-dense. There is often a roundish one (M_3_) at the base of the nucleus or at the base of the cell (M_4_). Nuclear variation is also very striking. Some nuclei (N_1_) are very electron-dense due to the abundant, dense heterochromatin. Other nuclei (N_2_) are less electron-dense. A third type of nucleus (N_3_) is homogeneous in appearance and intermediate in electron density. The fourth type nucleus (N_4_) appears to be the smallest and its heterochromatin slightly resembles that of the N_2_ nuclei. The fibroblast nuclei (FN) are somewhat different from the four types in the epithelium, but vaguely resemble the N_4_ type. **(b)** A pair of VS cells are unusually well preserved. Their vesicles (V) are numerous and apically located. They have few, short microvilli (Mv). One mitochondrion (M) is dividing, as is the one (M) in the cell portion to its right. Doorknob-like projections (DP) extend from a KS cell. JC, junctional complex. **(c)** A ciliated knob (K) of CR cell appears between two VS cells. The vesicles have a slightly greater electron density and the microvillus (Mv) is slightly longer than those in panel *A*. The structureless projection (SP) resembles the likewise rarely seen SP in the control 8 dpf larvae. All scale bars = 1 µm. KS = kinociliate support; CR = ciliated receptor; VS = vesicular support.

One KS cell (**Fig. 8a**) contained, in addition to the usual cilia and single type microvillus, a unique doubly-forked projection resulting in three branches, forming a microvillus group. The function of these microvilli (single, double or triple) is puzzling. The basal extensions from the KS cells (**Figs, 8a, 9a**) are suggestive of axons. The identification of two KS cells in addition to several VS cells and a MR cell with a forked microvillus (**Fig. 8b**) supports previous findings that single KS cells do occur removed from the groups that generally occupy the rims of the olfactory pits. In fortuitous sections where the plane of section and quality of preservation are both favorable (**Figs. 8c, 8d**), the vesicular and cytoplasmic membranes of a VS cell were ∼ 7 nm thick and consist of two electron-dense lines separated by an electron-lucent space.

**Fig. 8.**
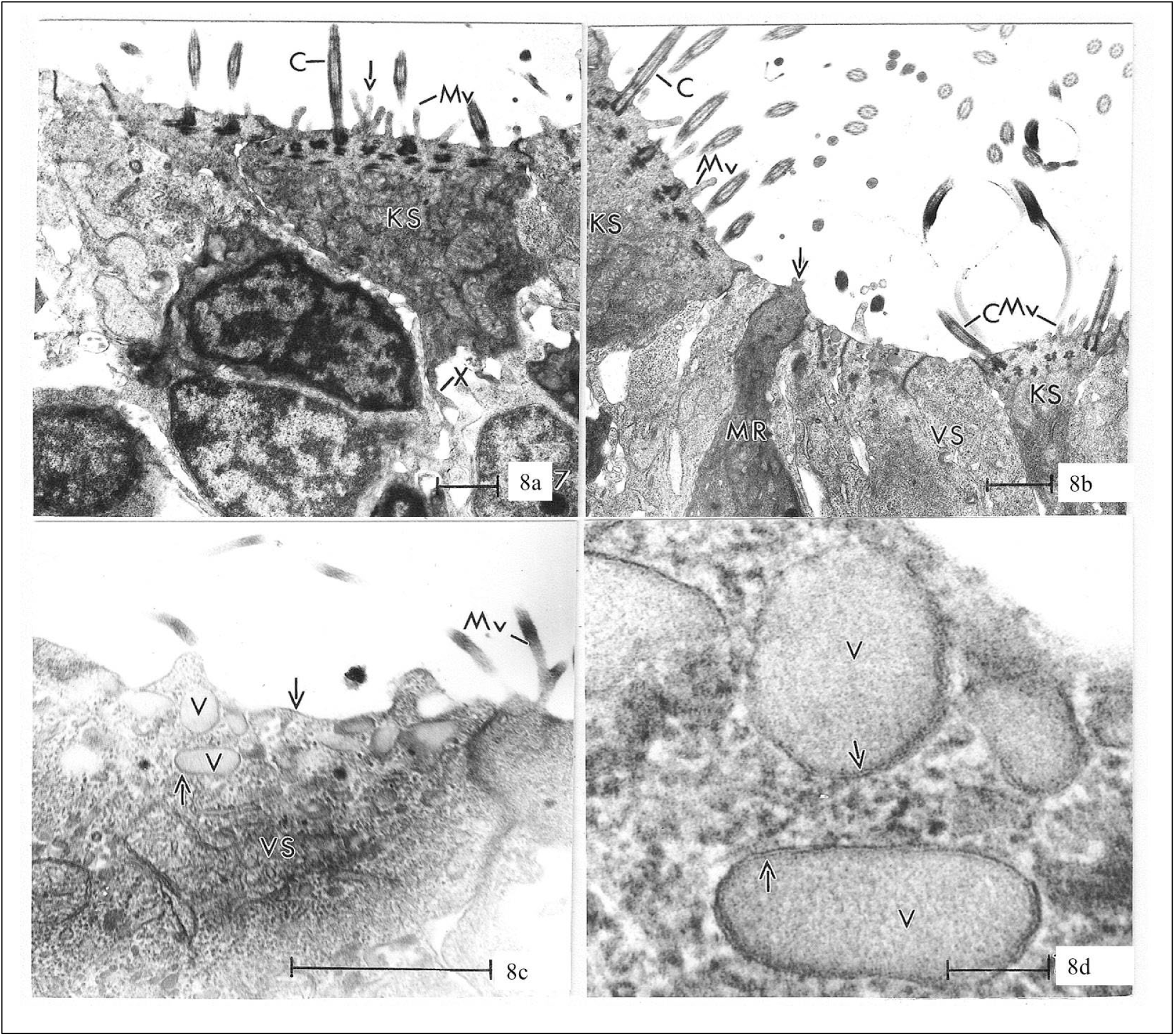
15 dpf control larvae. **(a)** The KS cell has both a simple microvillus (Mv) and a unique doubly-forked (arrow) microvillus group. C, cilium; X, presumed axon. **(b)** Two KS cells with cilia (C) and simple microvilli (Mv), a VS cell and a MR cell with a forked microvillus (arrow). **(c)** The VS cell has several vesicles (V) and vesicular and cytoplasmic membranes (arrows) that are about 7 nm thick and display a tripartite structure. Mv, forked microvillus. **(d)** A higher magnification micrograph of the tripartite structure of the vesicle (V) membranes (arrows). Scale bars in **(a)**, **(b)**, and **(c)** = 1 µm; Scale bar in **(d)** = 0.1 µm. KS = kinociliate support; MR = microvillar receptor; CR = ciliated receptor; VS = vesicular support.

In addition to the apparent axon extending from a KS cell, several presumed axons were observed (**Fig. 9a**). If these processes are axons, the cells are probably MR or CR cells. Both KS cells in the figure possess forked microvilli. A group of seven cilia, each with the same orientation of the central microtubule pair, were also present (**Fig. 9b**).

**Fig. 9.**
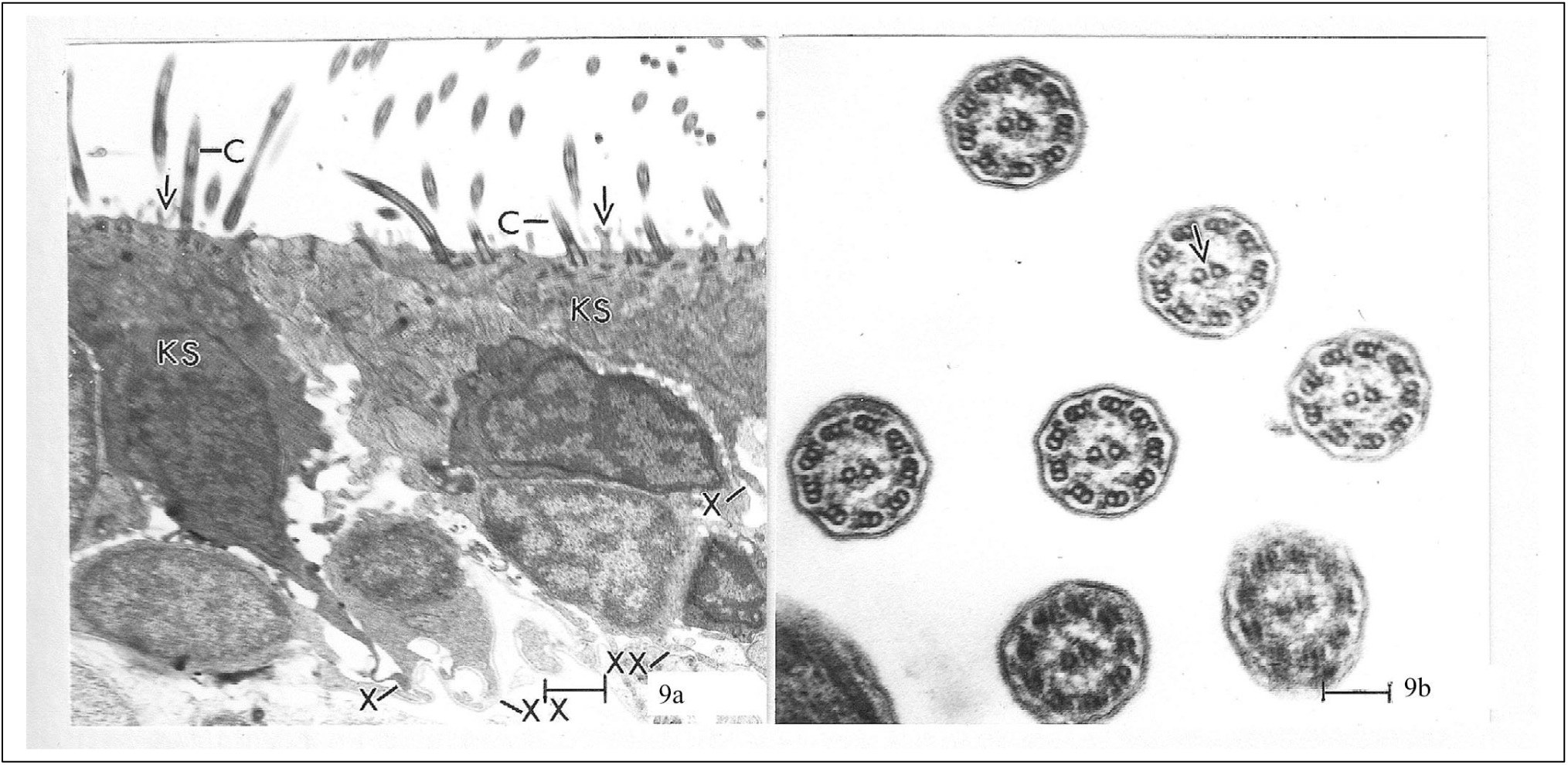
15 dpf control larvae. **(a)** The KS cells display cilia (C), forked microvilli (arrows) and an apparent axon (X). Non-KS cells (lower cytoplasmic and nuclear electron densities) also project what appear to be axons (XX). **(b)** This high magnification figure of cilia with identically oriented central microtubule pairs (arrow) illustrates the breadth of occurrence and the precision of orientation of this feature. Scale bar in **(a)** = 1 µm; scale bar in **(b)** = 0.1 µm. KS = kinocilitate support.

In sum, the olfactory epithelium at 15 dpf includes (1) forked microvilli projecting from both MR and KS cells, (2) doorknob-like processes from basally located cells, (3) nuclear variation, (4) mitochondrial variation, (5) vesicle structure variation, (6) vesicle number increase, (7) isolated KS cells near the center of an olfactory pit, (8) groups of preferentially-oriented cilia of KS cells, (9) a thinner olfactory epithelium, (10) doubly-forked microvilli, (11) axon bundles, and (12) a structureless projection (**Table 1**).

### Constant light: 4 and 8 dpf

To determine if constant light affects growth of the olfactory epithelium, we reared zebrafish to 4 and 8 dpf in constant light conditions prior to EM analysis. 15 dpf larvae are not included in this analysis due to poor survival in the constant light treatment group.

#### 4 dpf

In constant light-reared larvae, overall development of the olfactory system at 4 dpf is essentially identical to controls, with two olfactory pits, two eyes, forebrain, cartilage plates and mouth (**Fig. 10a**). Portions of three KS cells adjacent to the epidermal cell with its many projections and a nucleus are present (**Fig. 10b**). The olfactory epithelium is stratified and columnar. The basal lamina underlies the olfactory epithelium at the right edge of the figure. A section (**Fig. 10c, 10d**) cut parallel to the surface, showing the perimeter of the opening of the olfactory pit, reveals one of the only two mitotic figures found in this study. Surprisingly, the cell undergoing mitosis is positioned at the luminal edge of the pit and not basally (as in **Fig. 11a**). The presence of vesicles in the dividing cell suggests that it is a VS cell.

**Fig. 10.**
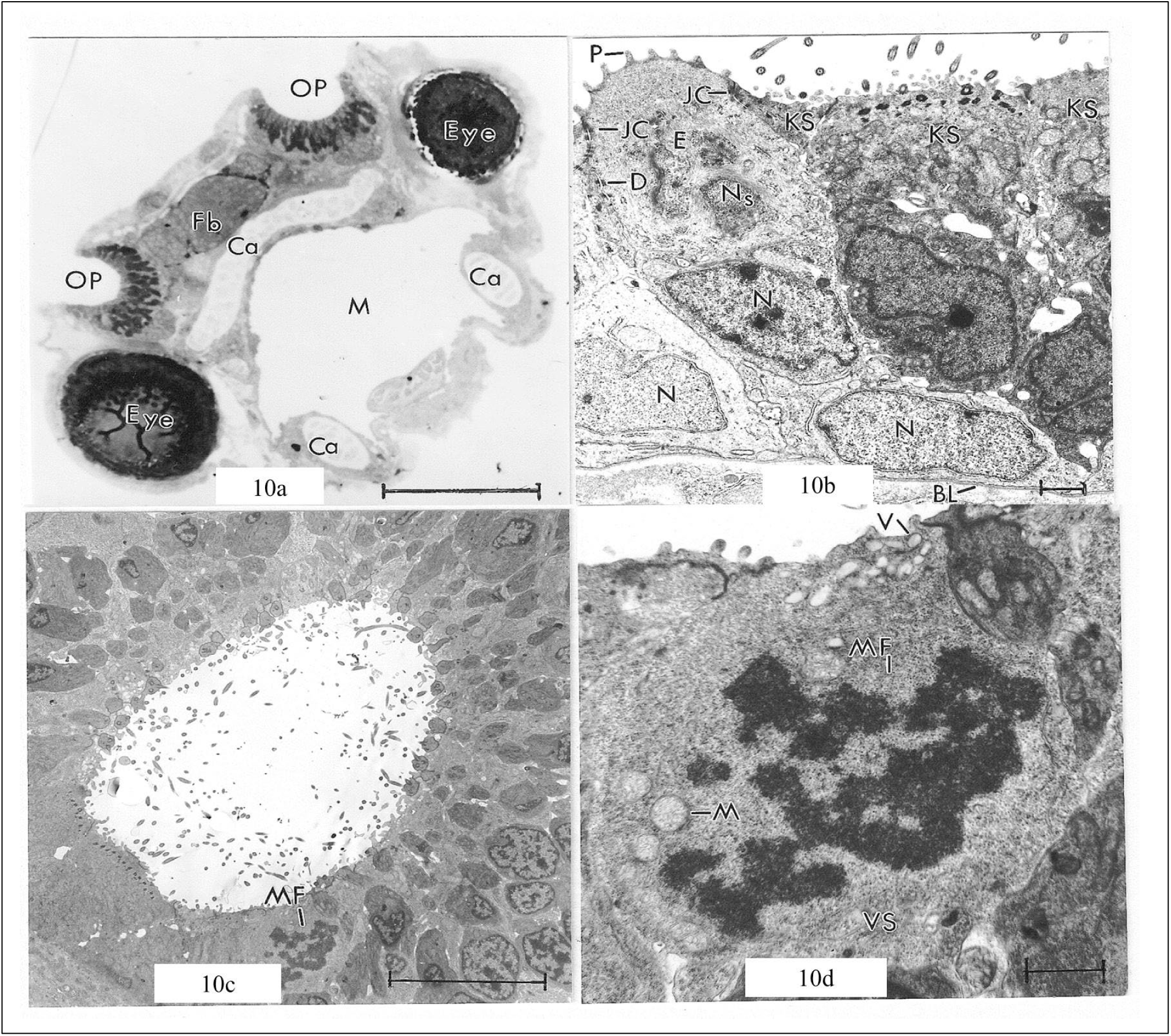
4 dpf constant light reared larvae. **(a)** Head of a 4 dpf larval zebrafish reared in constant light. The plane of section includes olfactory pits (OP), eyes (Eye), forebrain (Fb), several associated cartilage plates (Ca) and the mouth (M). **(b)** At this junction of the skin epidermis (E) with the KS cells of the olfactory pit, the surface epidermal cell shows its projections (P), junctional complexes (JC) with a KS cell to the right and another skin epidermal cell to the left, a desmosome connection (D) with another surface epidermal cell and a superficially sectioned nucleus (Ns). BL, basal lamina; N, nuclei of skin epidermal cells. **(c)** This section cut parallel to the surface of the olfactory pit opening shows one of the two mitotic figures (MF) encountered in this study. **(d)** A higher magnification of the cell with the mitotic figure (MF) is a VS cell. V, vesicle; M, mitochondrion. Scale bar in **(a)** and **(c)** = 100 µm; Scale bar in **(b)** and **(d)** = 1 µm. KS = kinociliate support; VS = vesicular support.

**Fig. 11.**
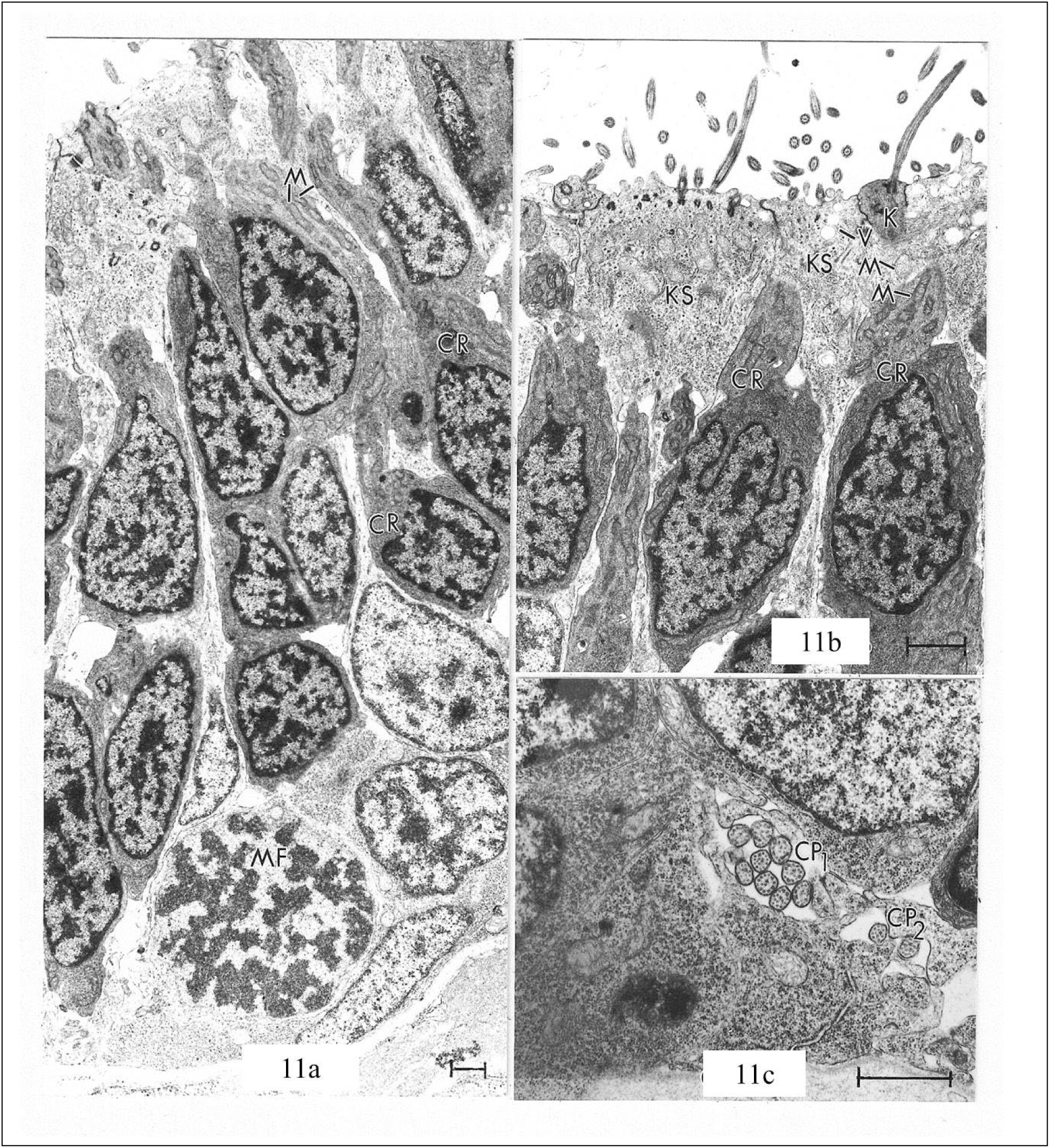
4 dpf constant light reared larvae. **(a)** The basal cell undergoing mitosis (MF) is the second of the two dividing cells encountered in this study. The first one was located apically in the olfactory pit epithelium; this one is located basally. The three nuclei aligned above the dividing cell indicate that this epithelium displays characteristics of a stratified columnar epithelium. The two CR cells to the right of the aligned threesome clearly illustrate the many elongated mitochondria (M) located in their dendritic extensions. **(b)** KS cells may contain vesicles (V). Their mitochondria (M) are more round and more electron-lucent than those of the CR cells. K, ciliated knob of ciliated receptor cell. **(c)** Two groups of circular profiles (CP_1_, CP_2_) appear basally in intercellular space. There is a marked variation in neurotubule arrangement among the different circular profiles. All scale bars = 1 µm. CR = ciliated receptor; KS = kinociliate support.

The three nuclei aligned in a row over the cell that is undergoing mitosis (**Fig. 11a**) further indicate a stratified columnar epithelium. The two CR cells to the right edge are an infrequently seen pair sectioned in almost exactly the same plane and showing that their long dendritic extensions are packed with mitochondria. Another section taken along a plane at ∼ 45° to the plane in Fig. 11a shows two similar cells (**Fig. 11b**) and also contains several broad-surfaced KS cells with vesicles. The shape and electron density of the mitochondria of KS cells are markedly different, i.e., they are rounder and less dense than the mitochondria of CR cells. Near the central portion (**Fig. 11c**) there is a group of nine circular profiles (CP) and an additional adjacent pair of circular profiles. Each circular profile contains neurotubules (microtubules). Notable is that each of the three best preserved profiles in CP_1_ contains a different array of neurotubules: one has 11 peripheral neurotubules plus 3 in the center; the second has 10 peripheral singlet neurotubules, one peripheral doublet and 3 central; the third has 7 peripheral singlet neurotubules, 2 peripheral doublets and 5 central singlets.

**Fig. 12a** provides a splendid opportunity to compare kinocilia and microvilli in a KS cell and to review the appearance of vesicles and CCR cells described earlier in 4 dpf control larvae. The microvilli are about two-thirds the diameter of the kinocilia. Incidentally, this KS cell is another example of how these cells are sometimes positioned a considerable distance from the rim of the olfactory pit. Vesicles are prominent in VS cells - one may be opening to the lumen of the olfactory pit. Several vesicles (bracketed in the upper right corner and enlarged in the inset) clearly show the presence of a limiting membrane and granules. In the deliberately more darkly printed inset (**Fig. 12a**, bottom right), they show a fine granular content. The CCR cell reveals two “in-sunk” cilia.

**Fig. 12.**
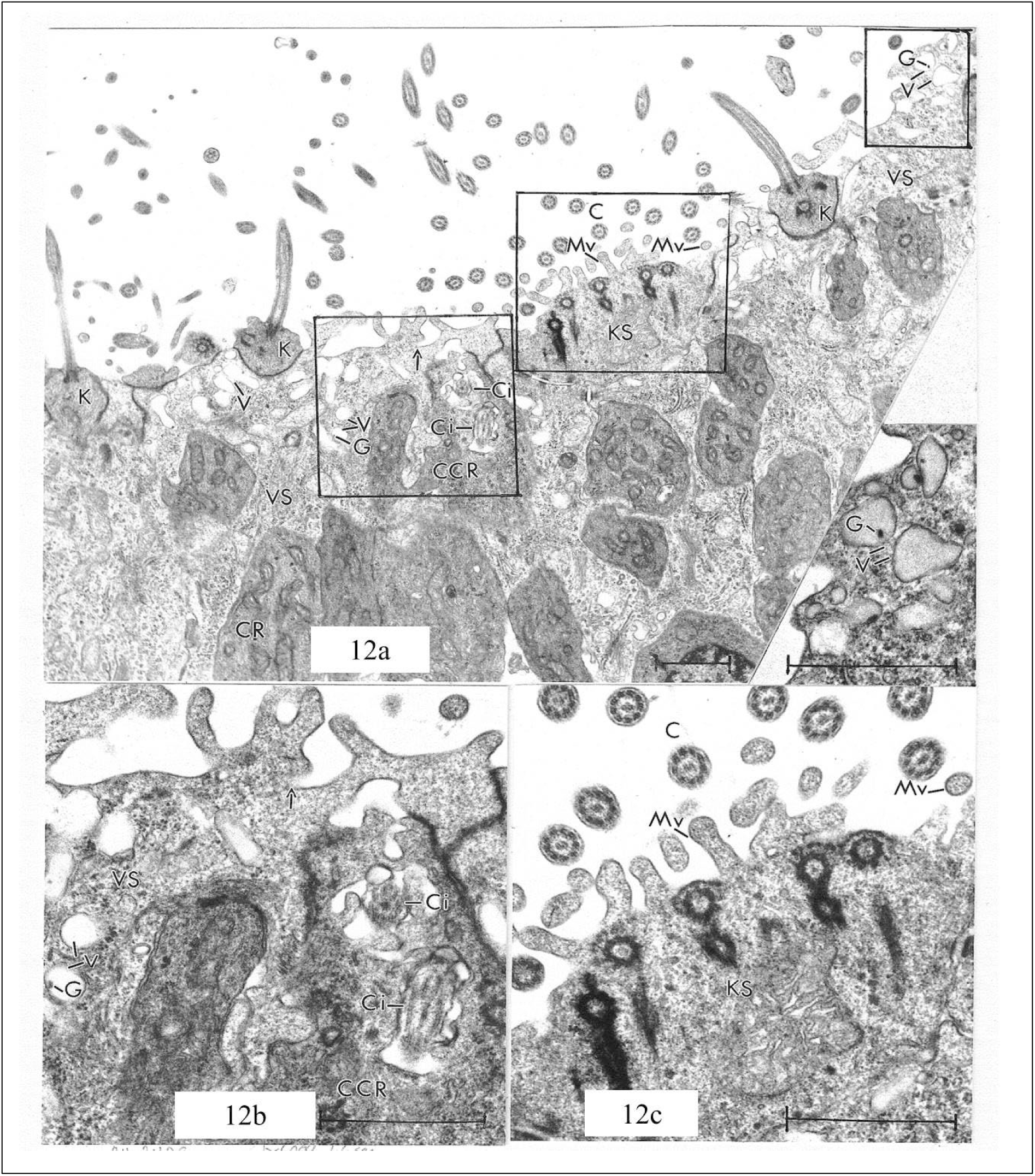
8 dpf constant light reared larvae. **(a)** Three ciliated knobs (K) of CR cells protrude from the surface. VS cells contain many vesicles (V), a few of which have an electron-dense granule (G), and one of which (arrow) appears to open to the lumen of the olfactory pit. A CCR cell with two “in-sunk” cilia (Ci) appears. The KS cell provides both longitudinally and transversely sectioned microvilli (Mv). The inset is an enlargement of the upper right bracketed area. **(b)** Enlargement of left bracketed area. **(c)** Enlargement of middle bracketed area. All scale bars = 1 µm. CR = ciliated receptor; VS = vesicular support; CCR = ciliated crypt receptor; KS = kinociliate support.

Thus, constant light rearing resulted in the identification of two mitotic figures at 4 dpf, one located basally and the other located apically within the olfactory epithelium. The apical mitotic cell appears to be a VS cell. Basally located groups of axons containing neurotubules with unusual paired relationships were also observed (**Table 1**).

#### 8 dpf

In general, ultrastructural components of 8 dpf constant light reared larvae were similar to those described above for control specimens of the same age. However, distinctly different dark and light basal cells within the olfactory epithelium (**Fig. 13**) were observed. These are most likely precursors to the labeled CR cell, which are dark, and the VS cells, which are light, located directly above them. We suggest this because of the density similarities and positions of the cells.

**Fig. 13.**
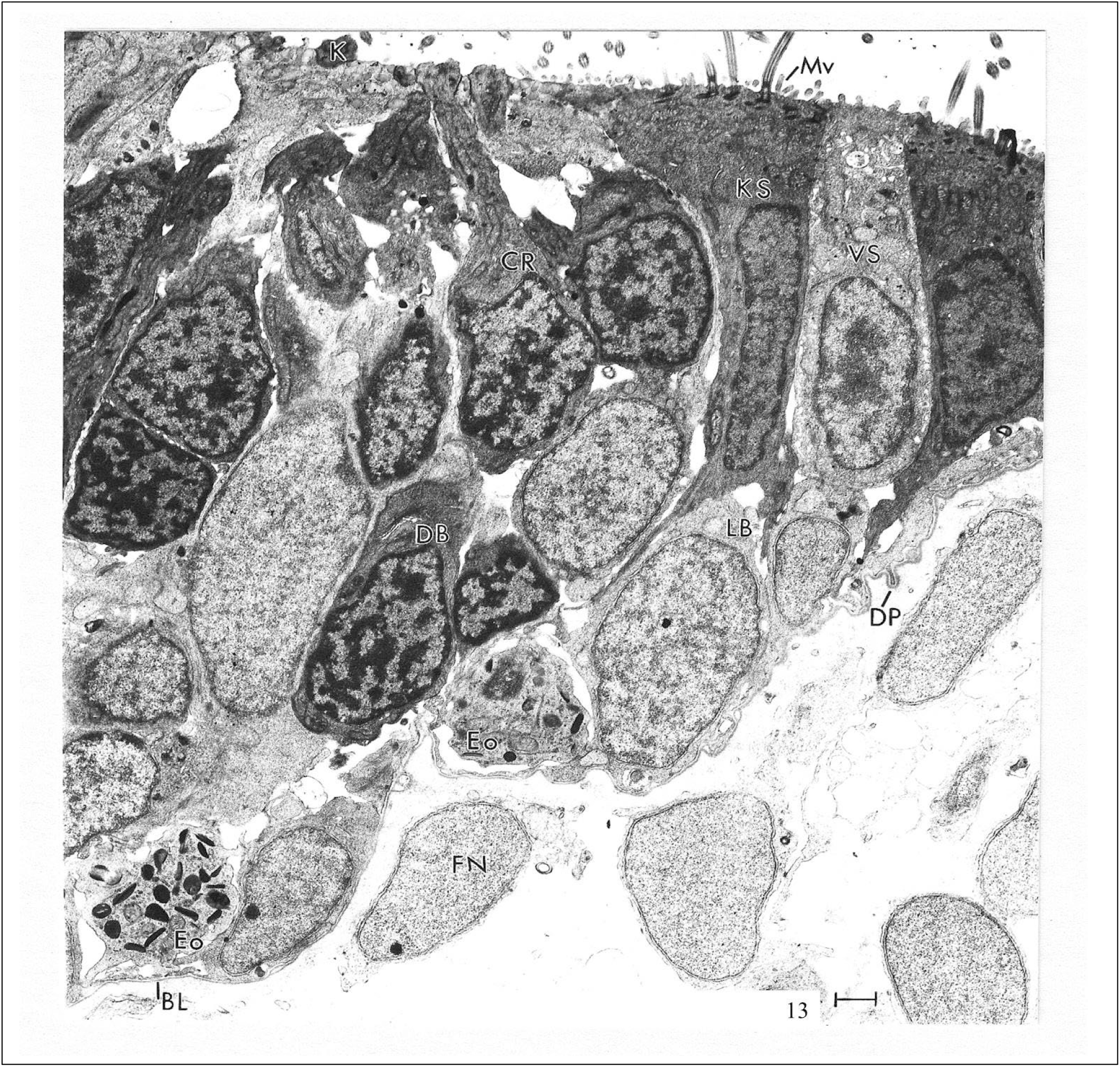
8 dpf constant light reared larvae. The figure permits a comparison of the appearance of the dark basal cells (DB) with the CR cells and of the light basal cells (LB) with the VS cells. The respective similarities in each case suggest that the dark basal cells are precursors of the ciliated receptor cells and that the light basal cells are precursors of the vesicular supporting cells. Possible precursors of the MR cells (not in figure) and KS cells are not so obvious. Portions of two neutrophils (Ne) appear in the basal region of the olfactory pit epithelium, as does a doorknob-like protrusion (DP). K, knob of ciliated receptor cell; Mv, microvilli of KS cell; BL, basal lamina; FN, fibroblast nucleus. All scale bars = 1 µm. CR = ciliated receptor; VS = vesicular support; MR = microvillar receptor

The most unusual feature of larvae reared to 8 dpf in constant light conditions was the occurrence of portions of two neutrophils in the basal region of the epithelium of the olfactory pit. A rarely encountered CCR cell was also found (**Fig. 14**; **Table 1**).

**Fig. 14.**
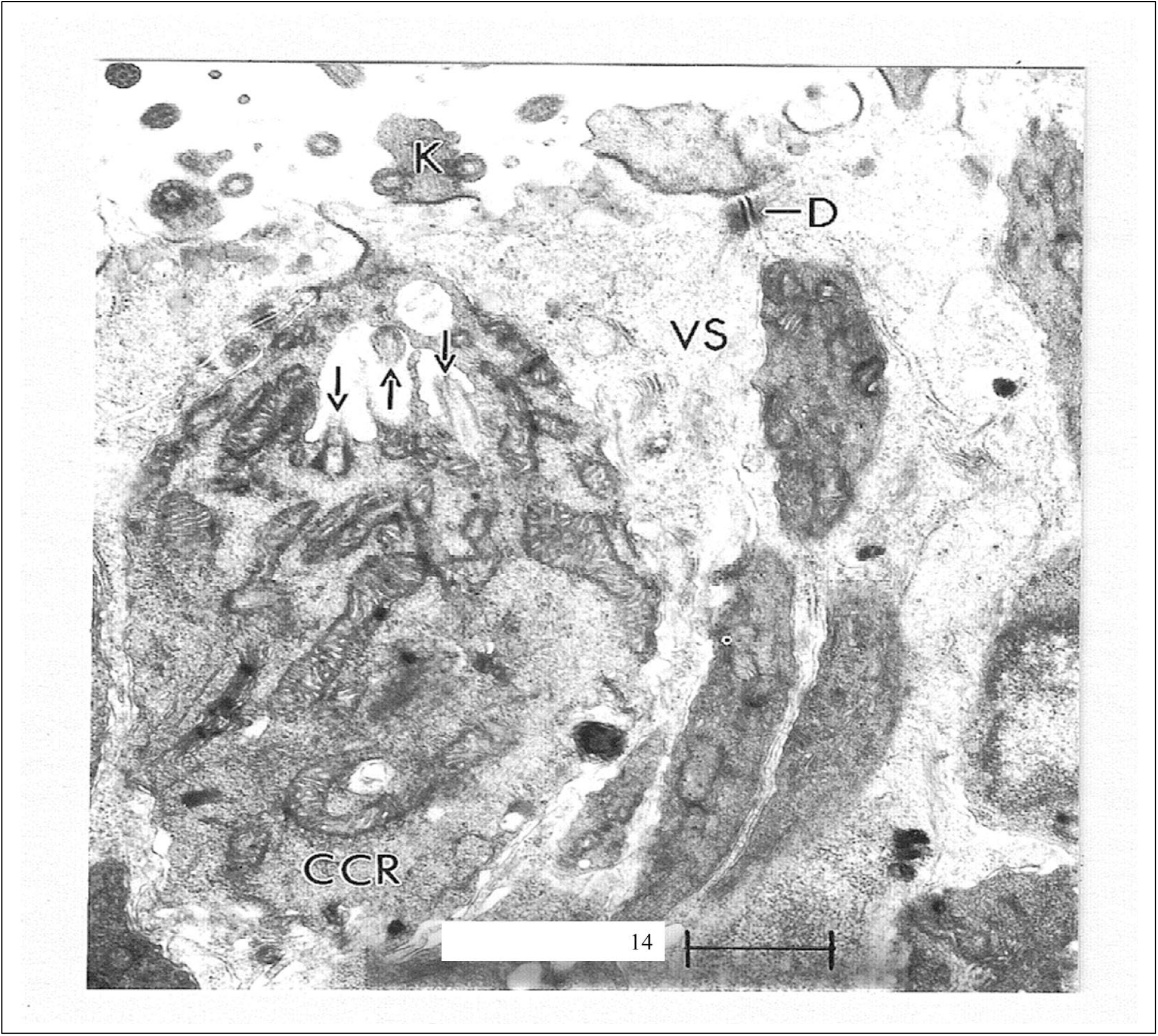
8 dpf constant light reared larvae. Micrograph of a ciliated crypt receptor cell (CCR) showing three “in-sunk” cilia (arrows) and many mitochondria (M). K, knob of CR cell; D, desmosome between two VS cells. Scale bar = 1 µm. CR = ciliated receptor; VS = vesicular support.

## Discussion

We report developmental changes in the larval zebrafish olfactory epithelium and how these structures are altered by constant light rearing. Age-related changes include centrally-located KS cells and forked microvilli from a MR cell that were observed at 8 and 15 dpf, but not at 4 dpf. In addition, several structures - axon bundles, individual axons from both a KS cell and a non-KS cell, microvilli from a VS cell, doubly-forked microvilli, doorknob-like projections, structureless projections, mitochondria of different electron densities, and the thinnest olfactory epithelium - were only observed in the oldest larvae examined (15 dpf). A monocilium was observed only at 8 dpf. These developmental changes suggest the olfactory epithelium continues to grow, including an increase in the number of receptor cells^5^ as the fish ages, with significant ultrastructural differences observed after 2 weeks of development.

### Olfactory epithelium development: 4, 8, 15 dpf

At 4 dpf, olfactory stimuli can reliably elicit behavioral responses^9,10^ and the olfactory pits have round openings and receptor cells are present.^5^ Our study indicates that CR cells are the dominant receptor type at this age; MR cells are not present and ciliated crypt receptor cells are rarely encountered. KS cells are primarily located around the periphery of the olfactory pit. VS cells are also evident, with active exocytosis of vesicle contents into the olfactory pit lumen. The vesicles in these larval cells bear little resemblance to either of the two vesicle types in the adult zebrafish,^6^ which were much larger in size, likely due to greater age. Cells in the olfactory organ of juvenile and adult garfish *Belone belone* are reported to synthesize and release a product stained by Alcian blue which stains acid mucopolysaccharides^43^ and vesicles within support cells were also identified by Hansen and Zeiske (1993). Neither cited report, however, illustrated a vesicle in the process of exocytosis, confirming release of substances into the lumen. The absence of any evidence of accumulation of vesicle content on the luminal surface of the pit may be due to the early age of the fish in this report or the removal of molecules once released.

The identified KS cells were located primarily around the rim of the olfactory pits. This preferential peripheral arrangement provides an example of how a structure can augment function at the cellular level. The peripheral orientation assures that incoming water will flow over the entire surface of the sensory epithelium of the olfactory pit. At the subcellular level, the preferential orientation of the central microtubule pairs in the ciliary axonemes implies that kinocilia movements direct water over/into the olfactory epithelium.^6^

At 8 dpf, all cell types are present in the olfactory epithelium – CR, MR, CCR, KS, and VS cells - and forked microvilli are now observed projecting from MR and KS cells. CR cells are still most abundant receptor cell type, greatly outnumbering MR cells. KS cells remain peripherally around the pit, an orientation that seems to disagree with the description of their distribution in adult zebrafish.^6^ Vesicles within VS cells are evident. Nucleosome densities vary among cells within the olfactory epithelium, suggesting speculation about which of the cell types gives rise to most of the more apically located cells, as its nucleus more closely resembles their nuclei than do the other two types of basal cell nuclei. Mitochondrial density also varies among cells. Mitochondria in the MR cell have a much greater electron density; while VS cells have the lowest cytoplasmic electron density and contain abundant but low electron dense mitochondria. It is possible that the low electron density of the mitochondria is related to the sparseness of the vesicles. However, mitochondria observed in these cells at 4 dpf were also low in electron density; perhaps they become low in electron density after the vesicles are formed and while inactive. The numerous large mitochondria in KS cells are reminiscent of those described in the supranuclear region of the kinociliate cells of embryonic and juvenile garfish *Belone belone*.^43^

The literature contains different classifications of the type of epithelium comprising the teleost olfactory mucosa. Bannister (1965) described it as simple columnar epithelium, while Byrd and Brunjes (1995) describe the zebrafish olfactory epithelium as pseudostratified, in agreement with others^43^ but contrary to Hansen and Zeiske (1998). In our micrographs, we identified many layers of cells between the underlying basal lamina and the free surface (**Fig.4b**) strongly suggesting a stratified columnar epithelium. However, if the CCR cell is, as accepted, a receptor cell, with no synapse in the olfactory epithelium, then it may be appropriate consider that the olfactory epithelium displays features of simple, pseudostratified *and* stratified epithelia. These differences could also depend on the age of the fish and the plane of the section. Finally, in regard to the epithelium of the olfactory pit, it should be recalled that only receptor cells need to extend from the free surface through the basal lamina. No synapses between receptor cells were observed in the epithelium.

At 15 dpf, there was an increase in the number and variety of cell types within the olfactory epithelium, consistent with continued growth and addition of cells. Presumed axons and axon bundles were also evident and differences in mitochondrial and nuclear densities were particularly apparent. For example, in KS cells, mitochondria were located supranuclearly and they were elongated and very electron dense like the cytoplasm of the cell. Mitochondria in VS cells, in contrast, tend to be similar in location and shape but lower in electron density and there is often a roundish mitochondrion located either at the base of the nucleus or at the very base of the cell.

### Constant light rearing: 4 and 8 dpf

In general, constant light conditions did not alter overall development of the olfactory system, though there were some structures that were observed only in this treatment condition. For example, in larvae reared to 4 dpf in constant light, mitotic figures were observed. Surprisingly, one of the only two mitotic figures found in this study was positioned at the luminal edge of the pit, and not basally as reported in adult zebrafish.^6^ This peripheral mitotic cell was identified as a VS cell, suggesting that the cell is either dividing after differentiating and migrating apically or that it delayed completion of a mitotic cycle that began basally while it migrated and differentiated. The latter explanation is supported by studies identifying a light-entrained circadian clock in developing zebrafish embryos and the importance of this oscillator in cell cycle progression.^44-46^ Specifically, it has been shown that constant light delays cell proliferation by suppressing several key regulators of mitosis that normally set the time for mitosis to late night/early morning.^45^ One consequence of this is the absence of a mitotic rhythm in cells exposed to continuous light conditions. One can speculate that perhaps the presence of mitotic figures only in the constant light condition in the current study is a result of a light-induced delay in mitosis to late morning (the time of euthanasia of the larvae). If true, that would also explain the absence of mitotic figures in the cyclic light/control conditions.

Continuous light exposure through 4 dpf also resulted in an absence of MR cells and the identification of KS cells within the olfactory pit (in addition to around the periphery). KS cells were not identified centrally until 8 dpf when larvae are reared in control conditions, suggesting early development of cells in this region as a result of constant light. Similar precocious development has also been shown for the visual system^27,28^ and accelerated growth is reported for zebrafish larvae reared in constant light.^24^

Animals reared in constant light until 8 dpf contained dark and light basal cells in the olfactory epithelium. These are most likely precursors to the cells directly above them, i.e., a CR cell (dark) and a VS cell (light). We suggest this because of the density similarities and positions of the cells. If this is the case, it raises the question of what cell type would be the precursor to both MR cells and the non-sensory KS cells. We do not have any ultrastructural data to support what these cells would be, though it is possible that the dark and/or the light basal cells are versatile in their ability to differentiate. MR cells were still absent, suggesting constant light conditions are deleterious to the development of this receptor cell type.

## Conclusions

Development of the zebrafish olfactory system begins early, with a defined placode at ∼17 hpf.^7,8^ Pioneer axons emerge from this placode at ∼24 hpf,^5,7^ which is about the time that the first olfactory receptor cells are born and olfactory receptor genes begin to be expressed.^13,14^ At hatching, CR and MR cells are present in the olfactory epithelium^5^ and odor responses can be detected in the olfactory bulb.^9^ Our results suggest that constant light rearing does not impact this general timeline of events, although ultrastructural modifications were observed in constant light exposed animals.

The zebrafish adult olfactory system has been examined using electron microscopy^4,6,47,48^ and one study has examined development at the ultrastructural level.^5^ The previous larval study provides detailed descriptions through 4 dpf – older ages are mentioned but not described in as much detail. Here we build on that study by providing additional details of the olfactory epithelium at 4, 8, and 15 dpf. These results could be used for future behavioral studies with known olfactory mutants, such as *laure*^17,19^ or with studies examining circadian and/or clock gene expression in the zebrafish olfactory bulb, as has been reported in mice^31^ and *Drosophila*.^30^

## Conflict of Interest

The authors declare that the research was conducted in the absence of any commercial or financial relationships that could be construed as a potential conflict of interest.

## Author Contributions

All authors had full access to all the data in the study and take responsibility for the integrity of the data and the accuracy of the data analysis. Study concept and design: GC, RT, VC. Acquisition of data: GC, RT. Analysis and interpretation of data: GC, RT, VC. Drafting of the manuscript: GC, RT, VC.

## Funding

This work was supported, in part, by a grant from the American University Mellon Fund (to VPC).

## Notes

### Competing Interest Statement

The authors have declared no competing interest.

